# *Preprint:* Triggered sequential viral-transduction from collagen-based scaffolds for tissue regeneration

**DOI:** 10.1101/2025.05.13.653729

**Authors:** John J. Amante, Bridget Twombly, Naaz Thotathil, Cathal J. Kearney

**Author notes:** **Corresponding Author:** Cathal J. Kearney. Life Science Laboratories S629, 240 Thatcher Road, Amherst, 01003.

## Abstract

Chronic wounds are a major healthcare issue that are recalcitrant to many traditional treatments. Increasingly, tissue engineering scaffolds are being developed and translated to promote their healing. To control signaling in the wound environment, gene therapy approaches are being explored, with adeno-associated virus (AAV) becoming increasingly popular. One critical challenge in chronic wound healing is that the wounds do not progress through the typical wound healing cascade, with signaling getting stuck in the inflammatory/immature tissue formation phase. This motivated us to develop a system capable of triggered sequential release of viral vectors to drive coordinated signaling. By housing this system within a collagen-glycosaminoglycan (GAG) scaffold, we aim to provide a proven extracellular matrix template as well as the correct signaling profile for closure of chronic wounds. Our system consists of two alginate pockets within the collagen-GAG scaffold, which we use to control the release of AAV. The first pocket allows diffusion of one AAV therapeutic and the second pocket can be ultrasound-triggered using low-frequency stimulation to release the second therapeutic. Initially, we developed and characterized the system using a reporter AAV. At our high AAV loading, we got sustained release and GFP expression in HEK293T cells over 9 days from our system in vitro, but lower loading had minimal transduction. When this lower group was triggered with ultrasound, cells were successfully transduced. Finally, we demonstrated sequential release of AAV encoding clinically-relevant genes for angiogenesis. This system has the potential for broad applicability as it can be readily adapted to mimic a range of biological pathways.

## Introduction

Chronic wounds are a major issue facing healthcare providers – an estimated 1 billion people globally are at risk of developing a chronic wound [1]. Estimates put the cost of chronic wound treatment at 3% of developed countries healthcare spending, highlighting the economic burden [2]. One major challenge with chronic wounds is their failure to progress through the normal wound healing cascade. In a healthy patient, a wound would progress through four stages: hemostasis (or coagulation) – the wound is closed off with clotting; inflammation – the wound is cleaned via the macrophages, neutrophils, and inflammatory mediators released; proliferation (or granulation tissue formation) – where cells begin to regrow into the wound; and matrix remodeling – where the new tissue strengthens over time [3]. Chronic wounds typically arrest their progress between inflammation and early tissue formation, leaving the wound in a permanently inflamed and compromised state. This is attributed to dysregulated signaling, which fails to drive transition through the various stages in these wounds.

Current therapeutic options include offloading, surgical debridement, and reconstruction surgery, which are either not fully efficacious or necessitate challenging procedures [4,5]. Tissue engineering has increasingly been applied to improve patient outcomes [6], with the two major focuses being explored for chronic wounds being growth factor delivery and biomaterial scaffolds [7–10]. Direct use of growth factors has shown promise to date. For example, growth factor delivery using vascular endothelial growth factor (VEGF) and platelet derived growth factor (PDGF) has been shown to help revascularize the wound bed, promoting regeneration in mice, and PDGF gel has had clinical success [11]. However, growth factors have short half-lives necessitating high doses; thus, an attractive alternative is viral transduction which would drive host cells to upregulate the desired growth factors locally at the wound site [12–14]. There are many viruses currently being investigated for use in clinical settings; however, adeno-associated virus (AAV), a single-strand DNA virus is a lead candidate [15]. While several viruses have been used for cells associated with wound healing (e.g., lentivirus has been used to improve wound healing in fibroblasts and adipose-derived stem cells [16,17]), lab-produced AAVs are considered safer than alternative viruses. This is due to the fact that they are generally considered as non-integrating and do not permanently edit cells, making them highly attractive for research and clinical translation[14,18–20]. Some recent studies, however, have found AAV can integrate into certain cells (i.e., liver) [21]. While there is some concerns with AAV integration driving oncogenesis evidence of this has only been observed in mice AAV integration causing cancer in humans is contested event with conflicting reports in the literature [22,23].

Biomaterial scaffolds provide a functional matrix to the wound, allowing cells to attach, migrate, proliferate, and deposit new matrix [24,25]. Choices for biomaterials scaffolds include chitosan, polyethylene glycol, and poly(lactic-co-glycolic acid) [26–28]. One commonly used type of biomaterial scaffold is collagen-glycosaminoglycan (collagen-GAG) scaffolds [29]. Initially developed in the 1980s, collagen-GAG scaffolds have seen clinical success over the ensuing decades and have been well characterized in the field [30,31]. Collagen-GAG loaded with growth factors have also been explored [32– 34]. While both growth factors and scaffolds are already being used in the clinic, no current clinical approaches consider the timing of delivery to reflect the body’s natural wound healing cascade.

As there is no universally successful approach to date, it is likely that these complex wounds need more complex and controlled solutions, for example mimicking the natural sequential presentation of growth factors/genes. Previously, wound healing in mice was greatly improved by following the timing in the angiogenic pathway as opposed to simply providing all growth factors at the same time [11,35]. Restarting angiogenesis has also been shown to restart the wound healing cascade, and revascularizing a wound improves patient outcomes [36,37]. Our lab has shown that simply aligning application of the VEGF and PDGF with their expression in healthy wound healing improved vasculature formation in vitro [38]. This data, combined with literature in the field, establishes motivation to design scaffolds for triggered release of therapeutics via ultrasonication of alginate hydrogels [7,39,40]. Our lab has shown that ultrasound can trigger therapeutic release from alginate hydrogels via cavitation-induced decrosslinking [39–41]. In previous work, the ionically crosslinked alginate of higher molecular weights (i.e., 250kDa) can be tuned for completely reversible decrosslinking/re-crosslinking when ultrasound is applied in physiological levels of calcium [41]. While reversible decrosslinking occurs with lower molecular weight alginates, some macroscale degradation is observed, which can be beneficial when delivering larger payloads, such as AAV as proposed here. While both timed growth factor delivery and collagen-GAG have each helped wound healing, the combination of both is less explored and should significantly improve patient outcomes. Unlike our previous work, in this system we propose using viral transduction to not just deliver a bolus of growth factor at a triggered time, but to switch on its production at a triggered time and then sustain protein expression throughout the wound healing cascade. We designed a collagen-GAG scaffold containing alginate pockets loaded with viral vectors encoding for various proteins (eGFP, mCherry, VEGF, and PDGF) and used diffusion or ultrasound-triggering to drive sequential release from alginate (see Figure 1). While some pockets will allow diffusion over time, others can be triggered to release at select times. By putting both pockets into one scaffold we can achieve sequential viral release and more closely follow biological pathways.

**Figure 1.**
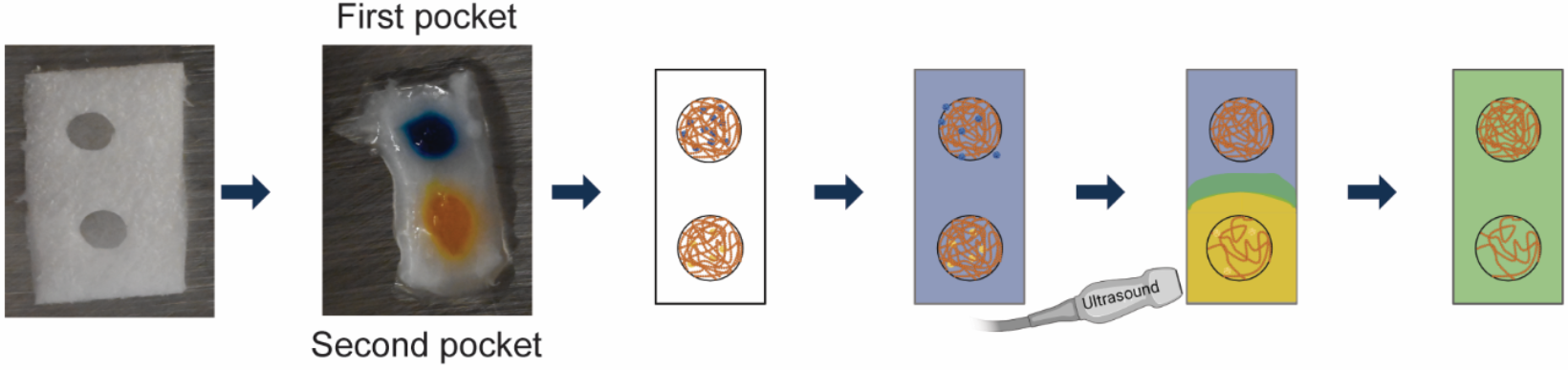
Schematic overview of the device. Starting with a collagen-GAG scaffold with pockets, alginate containing different adeno-associated virus (AAV) are injected into individual pockets. Higher dose AAV (modeled with blue) facilitates robust continuous diffusion, while the lower dose has limited diffusion. The lower dose (modeled with yellow) pocket can then be ultrasonicated to increase release at a user-defined time. This allows for sequential release of AAV to wound sites, within a proven regenerative template.

## Results and Discussion

### Alginate interferes with viral uptake

Before fully developing the release system, we first explored whether alginate itself affects AAV uptake by testing transduction in the presence of alginate. First, two different doses of AAV (either 1*10^9 gc or 1*10^10 gc of AAV-eGFP; no alginate) were added to HEK-293t cells and their signal read 2 days post-transduction via flow cytometry to confirm plasmid functionality. In both groups, close to 100% cells expressed eGFP (Figure 2A), thus confirming efficacy of the AAV-eGFP prior to its testing with alginate. To test whether alginate interfered with viral uptake, alginate, heparin (known to block AAV uptake), and chitosan (used as a negative control) were added to HEK-293t cells and then a multiplicity of infection (MOI) of 1 AAV-eGFP was added [39]. Interestingly, alginate reduced AAV uptake to a similar degree as heparin (Figure 2B). We also have verified that the AAV expression decreases over time, this data is included in Supplemental Figure 1. While alginate’s direct antiviral properties are being explored, the interaction between alginate and AAV has not been explored in detail [42]. Importantly, from the perspective of developing a delivery system, our data shows that there is a point where this interaction becomes saturated, such that an alginate gel loaded with a sufficient dose of AAV (e.g., 1*10^9 gc) will not inhibit transduction. This interaction – to the best of our knowledge – has not been reported in literature. Some groups have been exploring alginate as an anti-viral but to the best of our knowledge they have been more focused on viruses with a pathology [43]. When other groups have mixed AAV with alginate, the dosage was high enough so the interaction would have been saturated and remain unobserved [44]. Our data shows that AAV at a sufficient dosage could still transduce cells even in the presence of alginate but the observed phenomenon warrants further investigation.

**Figure 2:**
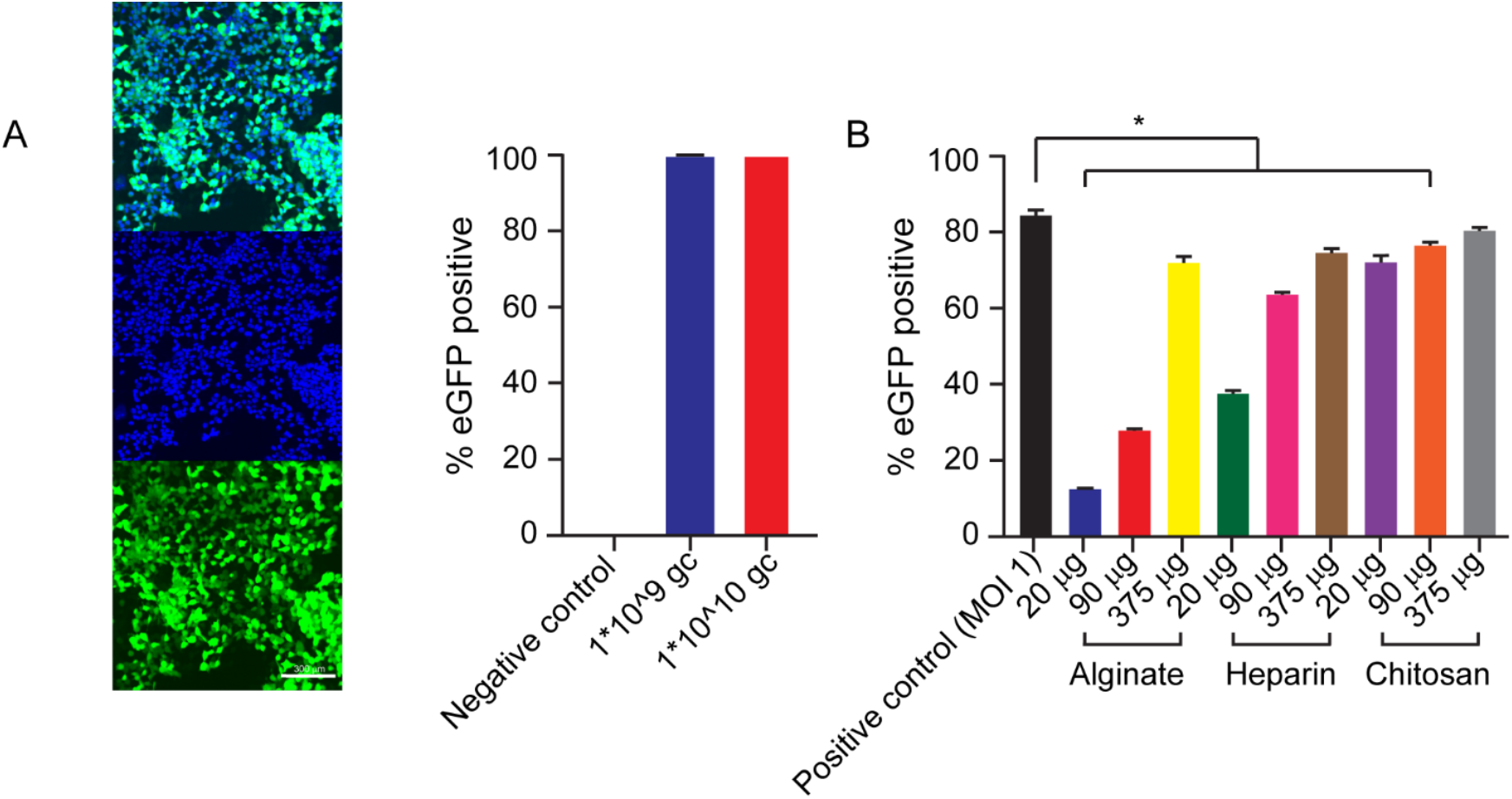
AAV interacts with alginate to reduce transduction. **A**. AAV-eGFP was added to HEK-293t cells at either 1×109 or 1×1010gc to confirm efficacy of our virus (left image: cells in well; right image, graph of flow cytometry data; n=3; error bars = mean ± st dev). **B**. AAV-eGFP (MOI = 1; 1.785*10^^^7 gc) was added with different concentrations of alginate, heparin, or chitosan polymer (non-crosslinked). The percent transduction was greatly reduced in the presence of mid to high concentrations of alginate and heparin, but not in the presence of chitosan (error bars = mean ± st dev; ANOVA with Tukey post hoc, * = p<0.05).

### AAV can diffuse from alginate gels and remain bioactive

Loading 1*10^9 gc/gel (low dose) or higher, we were able to bypass the alginate/AAV interaction and maintain a sustained release over 9 days (Figure 3A). We used AAV-eGFP and measured transduction, with GFP expression in flow cytometry used as a surrogate measure of AAV release and sustained bioactivity. Excluding the initial high percentage of expression over the first two days – attributed to an initial burst release of virus – overall transduction is remarkably stable over the course of the experiment, staying around 6% transduction per day showing diffusion happens over the full 9 days. A lower dose (1*10^8 gc/gel) does transduce but at barely detectable level over the same time frame (see Supplemental Figure 2A). Interestingly, transduction was not affected by alginate gel concentration as there was no statistically significant difference between a 1% and 2% gel (Figure 3A). Increasing the loaded dose to 1*10^10 gc/gel (‘high dose’) the transduction increased to an average of 85.8% per day on days 3 to 9 (Figure 3B). While the high dose maintained almost total transduction over 9 days the low dose stayed low enough that we anticipated that we could trigger increased expression by ultrasound, allowing us to sequentially deliver virus if loaded at the low dose. Prior to testing this we first confirmed the ability to transduce cells within a proven regenerative template.

**Figure 3:**
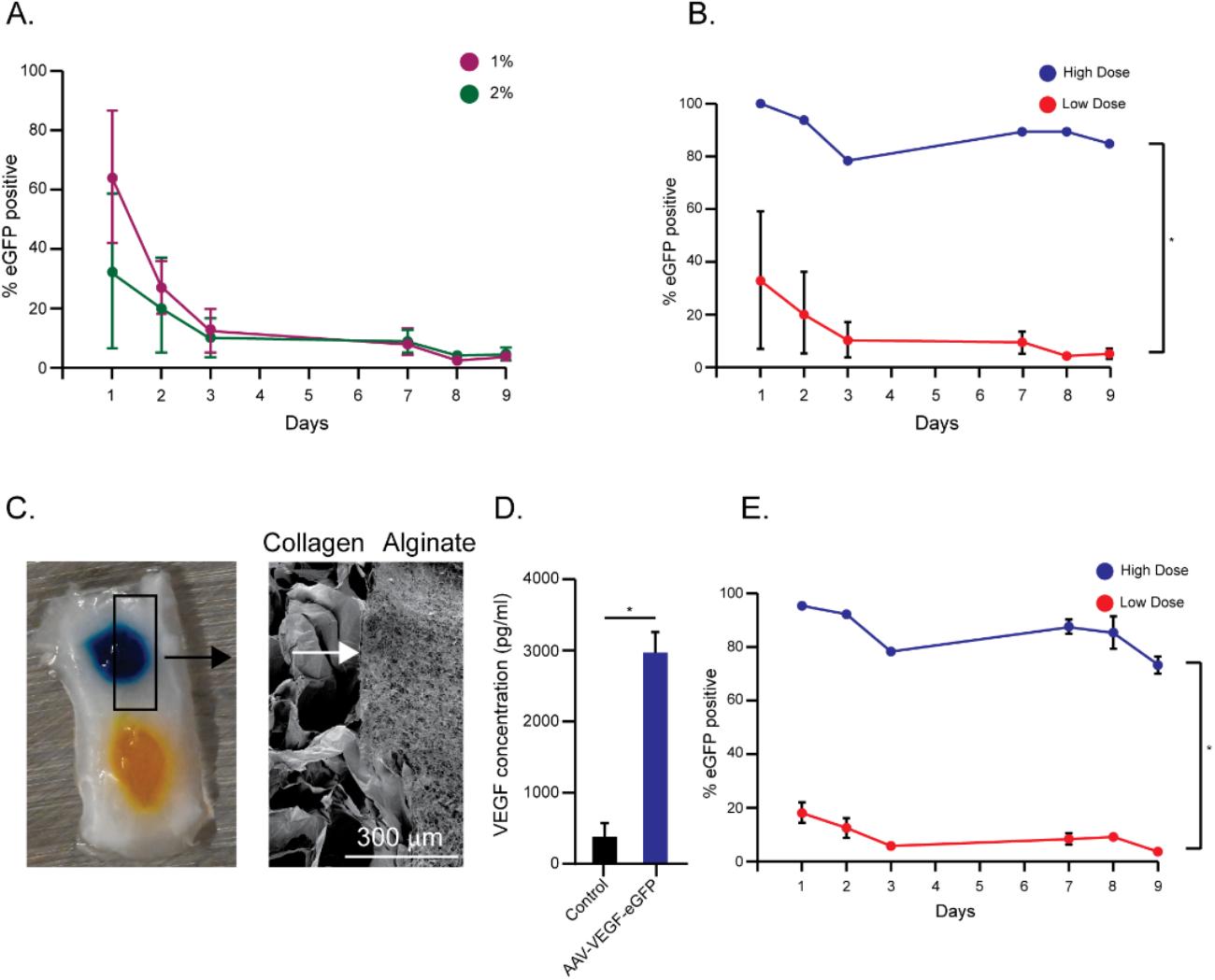
AAV successfully diffuse from alginate scaffolds. **A**. Crosslinked alginate gels at either 1 or 2% w/v were loaded with AAV-eGFP (low dose = 1*10^^^9 gc/gel) and allowed to diffuse over 9 days. While successful transduction was observed throughout the 9 days, no statistical difference was observed between gel concentrations. **B**. Alginate gels (1% w/v) were loaded with either low dose or high dose (1*10^^^10 gc/gel) and allowed to diffuse over 9 days and supernatant added to cells. At all timepoints, cells treated with the high dose had significantly higher %eGFP+ (error bars = mean ± st dev; t-test was used to compare at each timepoints, * = p=<0.05). **C**. Left: image of scaffold with alginate in pockets (food color added to alginate to make more visible) Right: SEM image showing the interface (white arrows) between alginate and collagen-GAG, confirming that alginate remains within the pocket. **D**. AAV-VEGF was added to coliagen-GAG scaffolds seeded with HEK293T cells. AAV-VEGF significantly increased VEGF output from the HEK293T cells. **E**. Alginate loaded with AAV-eGFP was injected into coliagen-GAG pockets and allowed to diffuse over 9 days and supernatant added to cells. While eGFP+ cells were observed in both groups, there was significantly higher %eGFP+ cells for the high dose group (Error bars = mean ± st dev, t-test was used to compare individual timepomts. * = p<0.05)

To see if this system could work within a collagen-based scaffold, alginate was injected into the pocket of a collagen-GAG scaffold and prepared for scanning electron microscopy to confirm alginate did not leak into the pores in the scaffold (see Figure 3C). Alginate naturally does not allow cell binding so we wanted to confirm the collagen-GAG scaffold would not be filled with alginate. The clearly demarcated interface between the alginate and collagen-GAG confirms that alginate stays within the pocket, giving us a pocket target that we can selectively ultrasonicate, and ensuring that alginate does not interfere with the regenerative portion of the scaffold.

With gels capable of viral transduction successfully developed, we next wanted to confirm that cells loaded on a collagen-GAG scaffold are capable of up-taking and expressing virus released from alginate. HEK-293t cells were seeded onto collagen-GAG scaffolds and AAV-VEGF (low dose) was added to the media. Conditioned media was collected 2 days later and an ELISA was run to check VEGF concentration. Viral transduction was confirmed as the AAV-VEGF exposed cells excreted more than a 7-fold increase in VEGF (Figure 3D). This showed that each individual component of the system could work, and next, we wanted to confirm functionality of the complete system (alginate loaded with AAV within collagen-GAG). A 1% alginate gel loaded with AAV-GFP (low and high dose run separately) was injected into a collagen-GAG pocket. Transduction was checked over 9 days (Figure 3E). Interestingly, incorporating alginate into collagen scaffolds eliminated the initial burst release. The low dose loaded scaffold averaged 9.8% transduction while the high dose averaged 85.6 %. With the diffusion pocket successfully developed, we began design and testing of the ultrasound-triggered release pocket.

### Ultrasound does not affect AAV bioactivity

First, to ensure ultrasound would not affect AAV bioactivity we ultrasonicated media containing an MOI of 1 AAV-eGFP for various times (60 seconds, 120 seconds, 180 seconds) either continuously or pulsed in 1 second intervals (Figure 4A). Ultrasound treatment times were picked based on temperature increases induced by ultrasonication, which we wanted to minimize to prevent AAV damage (Supplemental Figure 2B). Media was then added to HEK-293t and eGFP expression was compared to a non-ultrasound control and a negative control (no virus). We found that ultrasound up to 60 seconds does not statistically alter eGFP expression, showing that AAV can withstand some amount of ultrasound and maintain bioactivity. While it was not significant, we did notice some drop in bioactivity so to err on the side of caution we reduced the ultrasound duration for the remaining experiments to 30s.

**Figure 4:**
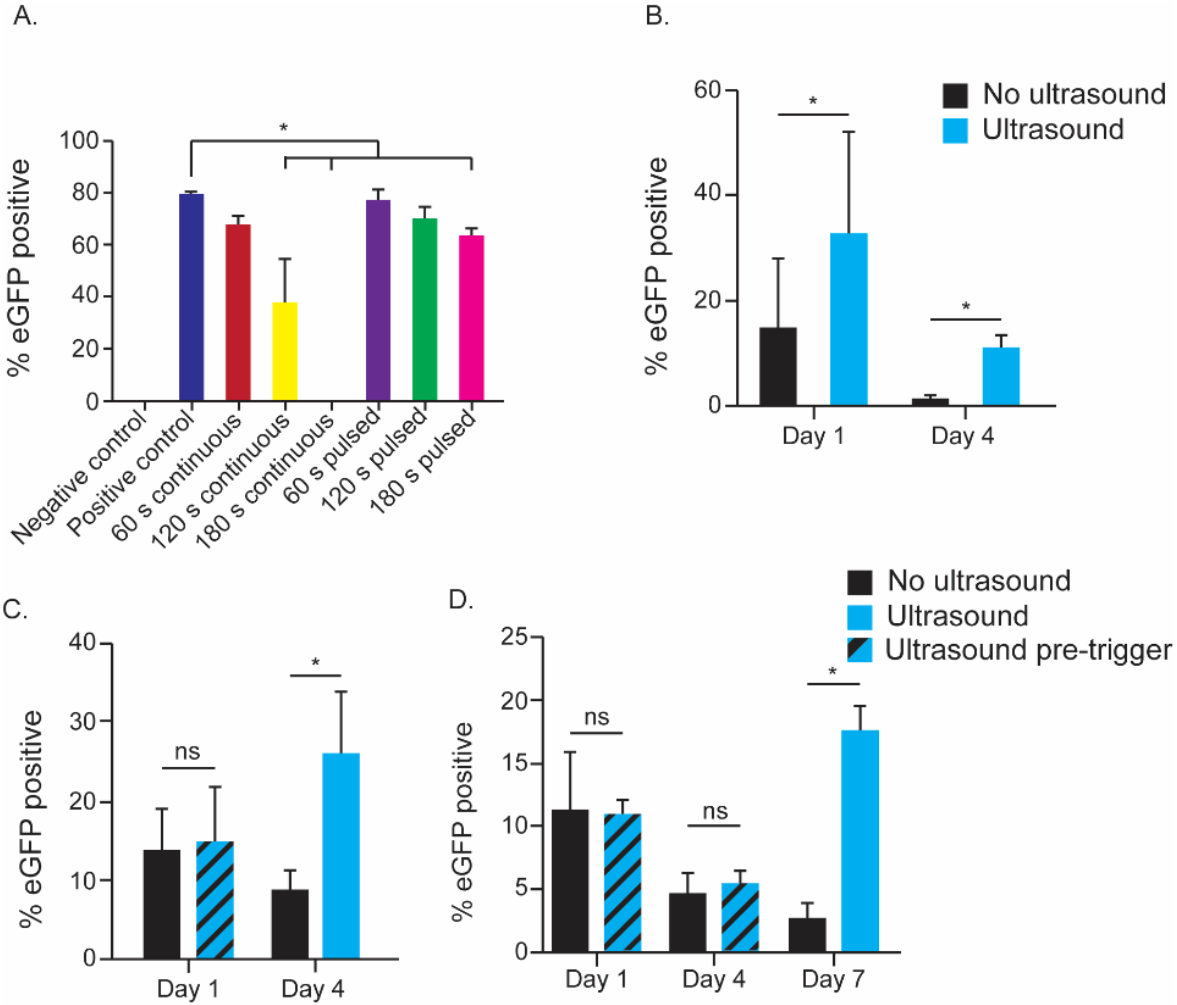
Ultrasound successfully triggers bioactivc AAV release. **A**. AAV-eGFP was suspended in cell media and ultrasonicated for various times either continuously or pulsed (1 second or/1 second off) Ultrasonication for 60 seconds did not significantly reduce bioactivity (error bars = mean ± st dev ANOVA with Tukey post hoc, * = p<0 05) **B**. Alginate gels batted with a tow dose of AAV-eGFP and injected into a coliagen-GAG scaffold were ultrasonicated on days 1 and 4 and compared to a non-ultrasonicated control Ultrasonication significantly increased % transduction on both days (error bars » mean ± st dev; t-test. * = p<0.05). C. Alg note gels loaded with a low dose of AAV-eGFP and injected into a coliagen-GAG scaffold were ultrasonicated only on day 4 The ultrasound group matches the control prior to ultrasound on day 1 but is significantly higher following ultrasound on day 4 (error bars = mean ± st dev; t-test, p=<0.05). **D**. Similarly, when the ultrasound group is left untreated until day 7, it matches control, but following ultrasound stimulation on day 7 it is significantly higher (error bars = mean ± st dev, 1-test, p=<0.05).

Alginate pockets loaded with a low dose of AAV-eGFP were ultrasonicated twice, on both days 1 and 4, and compared to pockets with equivalent low dose left untreated (i.e., no ultrasound; Figure 4B). On day 1 ultrasonicated pockets had a significantly increased amount of eGFP expression, with the ultrasound condition seeing a 2.7-fold increase. On day 4 this difference was more drastic with a 7.6-fold increase.

To further evaluate the potential of the system, we wanted to demonstrate different delayed-release profiles. First, we tested a diffusion phase followed by an ultrasound triggering on day 4. Alginate pockets loaded with low dose AAV-eGFP were measured for diffusion on day 1 and then with and without ultrasound on day 4 (Figure 4C). Both groups had equal diffusion on day 1; however, the ultrasound triggering increased expression from 8.8 % transduction to 26.1% on day 4. To further explore this, we allowed diffusion through day 7 in both groups and triggered the ultrasound group only on day 7 (Figure 4D). Again, on days without ultrasound the gels did not express different levels of eGFP; however, when ultrasonicated on day 7 the percent of transduced cells greatly increased from 2.8% to 17.7%. This demonstrates the ability to trigger release from these gels at user-defined times, allowing us to achieve sequential release. While we are not the first group to combine alginate and AAV, we believe we are the first to do so in a way that explicitly aims for both diffusion and triggered release in the same system. Several groups have looked at diffusion alone [44–46], while more recently groups have started exploring triggered AAV release [47,48]. Takatsuka et al. achieved their system’s release via near-Infrared light and alginate microbeads [47]. Ultrasound as a release trigger has been a focus of our lab, however we had not previously used it with a gene therapy approach [7,40]. While our system uses a different trigger, it fundamentally achieves the same goal of limiting AAV diffusion, and then allowing timed release. What truly sets ours apart is the ability to release more than one form of AAV encoding for different proteins from individual pockets. Other groups have started looking at dual release using a scaffold with two components allowing different diffusion rates [49,50]. The ability to define the trigger time offers a more flexible dimension to our approach than previously reported. One aspect of our research we were surprised to see – which was also observed by others – was the increase in transduction between days 3 and 7 when looking at diffusion alone; Lee et al noted a similar trend when working with fibrin glue [51]. This may be attributed to gel swelling dynamics. For example, monovalent ions replacing divalent cations in alginate gels and ‘weakening’ the gel; we previously saw this for ultrasound-responsive alginate; or it could be related to virus-alginate interaction dynamics [39].

### Sequential Release of Clinically-Relevant Genes

To demonstrate how our system could potentially be used in a sequential-release setup, we loaded two pockets with two different viruses within the same scaffold. One pocket contained a high dose of AAV-VEGF-eGFP, while the other contained a low dose of AAV-PDGF-mCherry (Figure 5A). These were chosen to mimic the vascular formation pathway whereas VEGF is expressed first to initiate vascular formation then PDGF is expressed to promote maturation of the vasculature [52]. Our logic was that the constant initial release of AAV-VEGF-eGFP loaded at a high dose would mirror the start of the angiogenic pathway, while the triggered release of AAV-PDGF-mCherry would mirror the later step of the angiogenic pathway. Both pockets were allowed to diffuse over three days and then on day 4 the PDGF-mCherry pocket was ultrasonicated. Using the fluorescent tags in the AAVs we checked plasmid expression over 4 days. eGFP expression between the no ultrasound and ultrasound scaffolds maintains close expression, whereas when ultrasound was applied, we see a large spike of mCherry expression. While the difference is consistent with our expectations, it is marginally insignificant (p=0.056)

**Figure 5:**
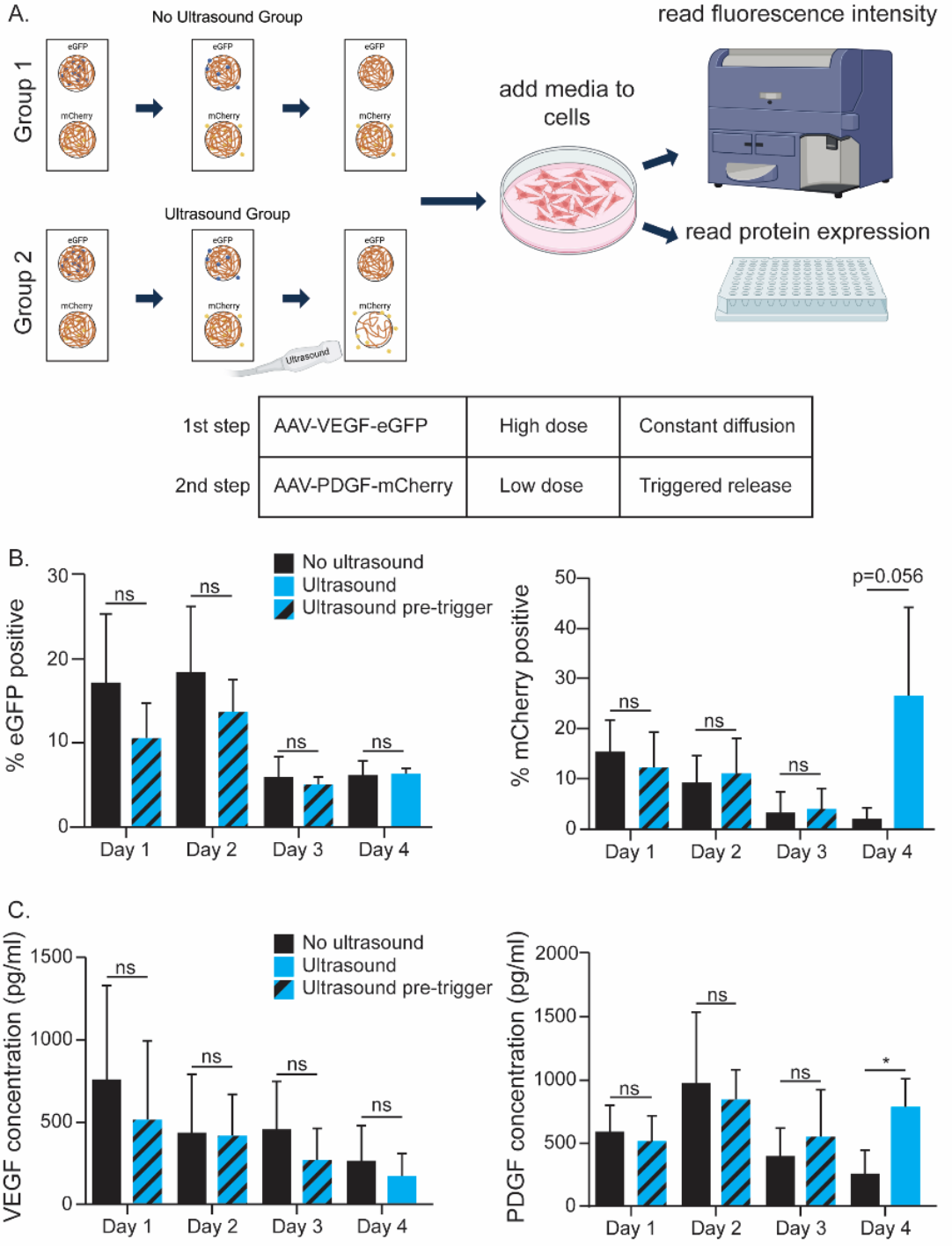
Incorporating clinically relevant genes into the device. **A**. Schematic overview of experimental setup The AAV-VEGF-eGFP is used to measure diffusion controlled release in the high dose group. AAV-PDGF-mCherry is used to monitor ultrasound-triggered release (day 4). Flew cytometry is used Io analyze the fluorescent expression, while ELISA was used to measure protein expression. **B**. Fluorescent expression e robustly observed tor eGFP and expression in the not ultrasonicated release pocket is not affected by ultrasound treatment of the otter pocket mCherry virus matches the no ultrasound group prior to ultrasound treatment on day 4. Triggering with ultrasound on day 4 enhances %mCherry+ cells (error bars = mean ± st dev). **C**. ELlSAs were run on the collected conditioned media. While VEGF expression was never significantly different between groups, the ultrasonicated PDGF conation was significantly higher than the diffusion control following ultrasound on day 4 (error oars = mean ± st dev: t-test. p=<0 05).

Conditioned media was collected each day and then ELISA was run to check both VEGF and PDGF protein levels in the conditioned media. Over the four days, VEGF expression (caused by the high dose of AAV-VEGF-eGFP) was never significantly different between both scaffolds, which shows that individual pockets can be triggered without affecting neighboring pocket release. By contrast, on day 4 the cells exposed to the ultrasonicated releasate from the low dose of AAV-PDGF-mCherry pocket expressed significantly different PDGF compared to the diffusion alone (Figure 5C). This confirmed our proposed approach that we could trigger AAV release from alginate using ultrasound while keeping a consistent diffusion release.

While the imaging and protein data show significant differences between ultrasound and diffusion only, we would like to further optimize the ultrasound so that the PDGF dose following ultrasonication is the maximum out of all the doses. Ultimately a cleaner difference would be ideal but by simply swapping out viruses without any additional optimization we have already achieved significance, showing that this system works and is highly adaptable. With a functional payload our system is set to move into preclinical testing with minor modifications, mainly reducing the thickness of the scaffold to be a similar thickness to mouse skin (roughly 2 mm). Otherwise, ultrasound has been shown to be safe to use across a wide range of functions and the device will be implanted in the patient’s skin meaning it will be easy to reach and be directly ultrasonicated [53–55]. Relevant to this application, a similar ultrasound profile was already tested in mice, and sonophoresis has been explored for transdermal drug delivery using similar ultrasound parameters [41,56]. Going forward we expect groups to have to adjust the titers of virus used in each pocket for their own goals. In our system we are not concerned with toxicity as AAV release is localized, reducing the risk of body wide adverse effects. Since each cell type does not uptake AAV the same, switching models might require minor adjustments to the amount of AAV loaded. We do not have concerns about overloading as while high dose of AAV has had adverse effects, they have only been noted in exceptional cases [57]. While AAV is generally considered non-integrating, some recent studies, however, have found AAV can integrate into certain cells and this potential should be considered prior to translation [21]. The authors note that low levels of integration were observed and suggest that cells with integrations will be replaced over time.

In the future, we plan on adding and testing additional clinically relevant genes. Further, as we were developing the system in this paper, we used a readily available cell line that is responsive to AAV transduction and we will transition to more clinically relevant cells (e.g., fibroblasts for wound healing). We did notice a reduction in cell growth when cells were combined with ultrasonicated media (see Supplemental Figure 2C), which needs further exploration also. It is possible that scaffold degradation products (e.g., crosslinker) are being released during ultrasonication and are detrimental to cells.

## Conclusion

In this report we determine how to achieve sequential release of AAV from a collagen-GAG scaffold. First, we identified how to achieve different release profiles, and then incorporated our gels into scaffolds, further characterizing the release. Finally, we demonstrated using ultrasound that we could trigger release of the AAV from our system at user-defined times, and demonstrated time-controlled release of VEGF and AAV. We are excited for the possibilities this will open up, as our system is readily modifiable to be used in a wide variety of applications.

## Materials and Methods

### Cell culture

HEK-293t cells were cultured in DMEM high glucose (Genesse Scientific, 25-500) with 10 % fetal bovine serum (Biowest, S1620) and 1% penicillin/streptomycin (Gibco 15140-122). Cells were left in a HeraCell VIOS incubator set at 37 degrees C when not actively being handled (CO2 set at 5%). When needed, cells were lifted with the following trypsin treatment for 2 mins (Quality Biological, 118-087-721; 1mL for T75 flask; 200 ml/well for 24-well plate). The default cell seeding density was 30k cells/well on a 24 well plate.

### Adeno-associated virus

AAV was purchased from Vector Builder and University of Massachusetts Chan Medical School Viral Vector core. All AAV were serotype 2 and encoded either eGFP, VEGF-eGFP, or PDGF-mCherry. To confirm our cells could be transduced, AAV at both 1*10^9 genomic copies (gc) and 1*10^10 gc was added to cells and uptake was confirmed as described using a flourescent microscope (EVOS) and flow cytometry.

### Flow Cytometry

To prepare cells for flow cytometry, samples were lifted with 200 µl of trypsin, spun down at 750 rcf, and resuspended in 450 µl of PBS before being put through a 0.22µm filter. Data was analyzed using FlowJo. This process was used in all tests done with flow cytometry. Transduction was read through flow cytometry on a LSRFortessa using the 488 nm laser (eGFP emission: 488 nm) (University of Massachusetts Amherst Flow Cytometry Core).

### Binding experiments

30k HEK-293t cells were plated and alginate, heparin, or chitosan solutions were added to 24-well plates in increasing doses. All solutions were 2% w/v and the volume added to wells, containing 500 ml of media, was adjusted so that the following quantities of polymer were added: 20 µg, 90 µg, or 375 µg (i.e., 1. µl, 4.5 µl, or 18.75 µl of 2% w/v solution). Next, a MOI of 1 AAV-eGFP (1.785*10^7 gc) was added to each well; the amount of AAV was constant for all samples. To prepare cells for flow cytometry, samples were lifted with 200 ml of trypsin, spun down at 750 rcf, and resuspended in 450 ml of PBS before being put through a 0.22µm filter.

### Gel formation

Alginate (from Novamatrix, UP LVG), phosphate buffered solution (PBS), and AAV were mixed between two Leuer lock-connected syringes. The mixture was then left for 1 hour before crosslinking by being mixed with an additional syringe containing PBS and calcium sulphate solution (final concentrations: alginate 10 – 20 mg/ml; 50mM calcium). Gels were then cast between two Teflon sheets with spacers, and allowed to crosslink for 30 minutes, and then moved into 500 µl of cell media (in 24 well plate). All gels were either 1% w/v or 2% w/v alginate. Gels were stored at 37°C and 5% CO2 in a HeraCell VIOS incubator. To test that AAV maintained bioactivity when put into the hydrogels, conditioned media was collected and added to a well containing 30k HEK-293t cells on each day recorded, and fluorescence was checked via flow cytometry using the method described in Binding Experiment. An example of the flow analysis is provided in Supplemental Figure 3.

### Scaffold fabrication

Based on previously described protocols, [10], collagen (Collagen Solutions) was soaked overnight in 0.05 M acetic acid at a concentration of 4.5 mg/ml. Then collagen was blended with an IKA Ultra-Turrex for 90 minutes. Chondroitin-sulfate 6 (Toronto Research Chemicals Inc.) that had been resuspended in 0.05 M acetic acid at a concentration of 2.66 mg/ml was added dropwise while the blender was running, and the solution was blended an additional 90 minutes. Collagen-GAG solution was then degassed and freeze-dried in a Christ Epsilon 2-4 freeze dryer at -10°C using stainless steel molds. To create pockets a negative mold with metal pillars was used.

### Inserting gel into scaffolds

The protocol for gel formation and scaffold fabrication was followed with only minor edits. Instead of casting gels onto Teflon, a 20 ½ gauge needle was connected to the syringe and the gels were cast into pockets in the scaffold. Release was still tested using release-conditioned media. Scaffolds were stored in 2 ml of media on a 12 well plate at 37°C and 5% CO2. A schematic of the work-flow is provided in Supplemental Figure 4.

### SEM

SEM was performed by Electron Microscopy Core Facility at the University of Massachusetts Amherst. Scaffolds were dehydrated via freeze drying (−10 °C over 24 hours at vacuum). Scaffolds were then sputter coated with gold and then imaged with an FEI Magellan 400 XHR-SEM. The acceleration velocity was 1 kV and the beam current was 13 pA.

### Triggered release gel in scaffolds

After inserting gel into scaffold, a Sonics Vibra-cell (model VCX130) was used to ultrasonicate the alginate in scaffolds. The ultrasound probe was inserted roughly 2 mm into the media and positioned above the targeted gel (without touching the gel itself). Viral release was triggered using 30% amplitude for 30 seconds unless other durations are specified in the figure caption. Afterwards, media was filtered through a 100 µm filter to clear loose alginate or collagen-GAG, and then added to cells. When scaffolds were tested on multiple days, they were stored at 37°C and 5% CO2. Cells were not collected after flow cytometry and each time point used its own cells.

### Dual Release experiment

Two different alginate gels were prepared, one at a high dose (1*10^10 genomic copies/gel) containing AAV-VEGF-eGFP and another at a low dose (1*10^9 genomic copies/gel) containing AAV-PDGF-mCherry. Both gels were inserted into the same collagen-GAG scaffold which contained two different pockets. Conditioned media was added to 30,000 HEK-293t cells on days 1,2,3, and 4. On day 4 the PDGF pocket was ultrasonicated. Between treatments scaffolds were held at 37° C and 5% CO2. Cells were analyzed for fluorescent expression with a Cytek Aurora flow cytometer using the same method described in Binding Experiment and the releasate was analyzed using ELISA (mCherry emission: 610 nm).

### ELISA

ELISA kits were purchased from R&D Systems. Instructions provided were followed for both kits. Kits were Human VEGF (catalog DY293B) and Human PDGF-BB (catalog DBB00).

### Statistics

Data was entered and analyzed in GraphPad Prism and graphs were made. For studies with two groups (e.g., ultrasound versus diffusion), paired t-tests were used as each gel was fabricated and then split into an ultrasound or diffusion group. If gels were treated over multiple days, t-tests were run to compare the two groups at each individual day. If there is more than one group, one-way ANOVA with Tukey’s post-hoc testing was used. When graphed, error bars are standard deviation as calculated by the software. Significance was defined as p<0.05 and indicated by * on graphs.

## Author contributions

John Amante: Conceptualization, data curation, formal analysis, investigation, methodology, project administration, supervision, validation, visualization, writing. Bridget Twombly: investigation. Naaz Thotahil: Investigation. Cathal Kearney: Conceptualization, funding acquisition, methodology, project administration, resources, supervision, writing

## Data availability

The data will be deposited into the Data Repository @ ScholarWorks following completion of peer review and the DOI provided

## Acknowledgements

The authors would like to thank the University of Massachusetts Amherst Flow Cytometry Core and the Scanning Electron Microscopy Core for their help and letting us use their equipment. Biorender was used to create schematics. Research reported in this publication was supported by the National Institute of General Medical Sciences of the National Institutes of Health under award number R35GM147272.

