## Supplemental Data for "*Preprint:* Triggered sequential viral-transduction from collagen-based scaffolds for tissue regeneration"

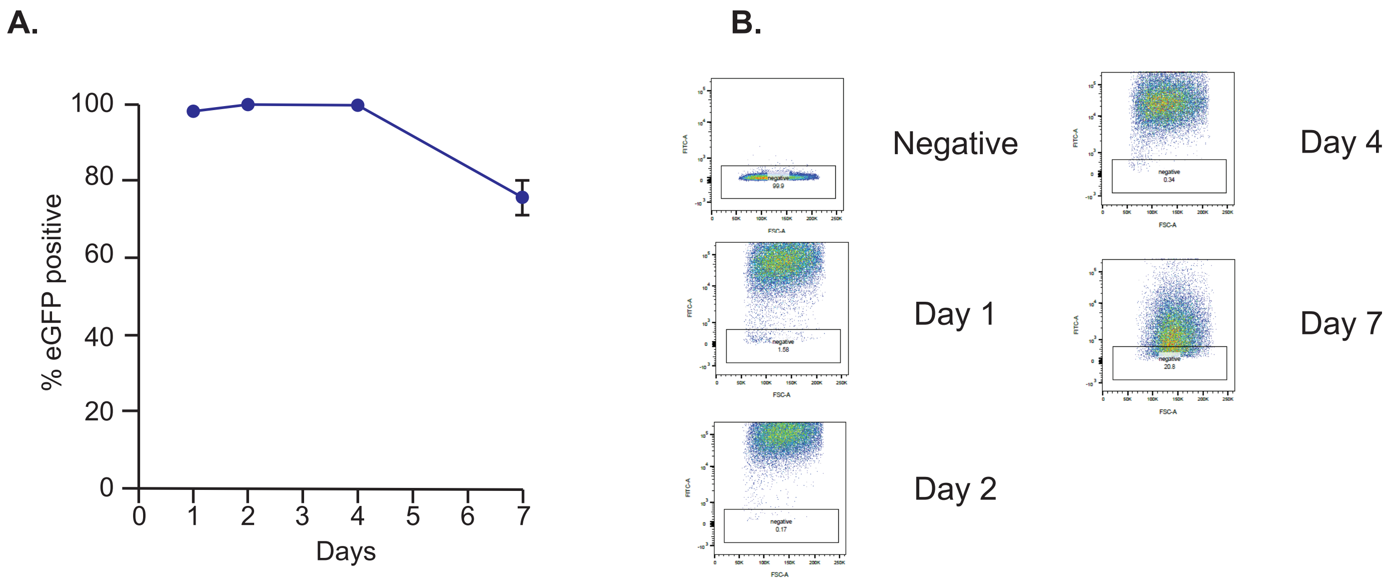


**Supplemental Figure 1: AAV does not significantly integrate into the cells. A.** Percentage of HEK293t cells expressing eGFP over time after addition of MOI 1 AAV-eGFP. The total amount of cells expressing eGFP decreases overtime, dropping down to 76 percent by day 7. **B.** Representative plots showing the overall intensity of eGFP over time. By day 7 it is clear the intensity as well as the number of positive cells are decreasing.

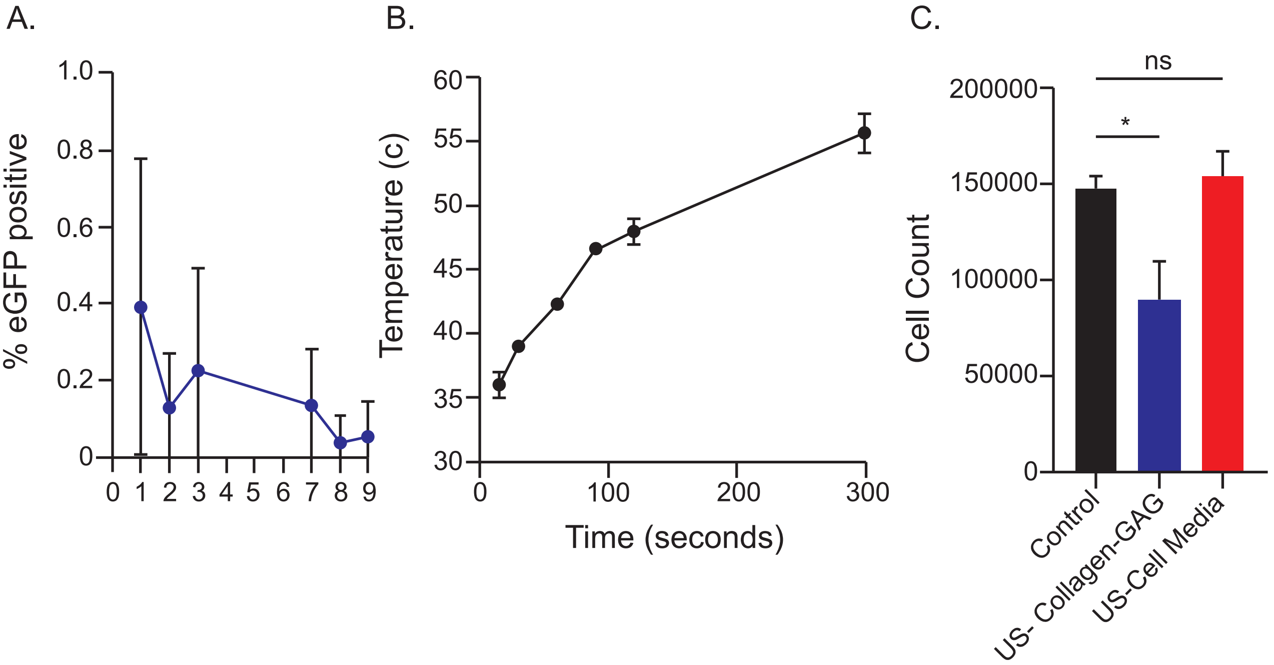


**Supplemental Figure 2. A.** 1*10^8 gc/gel of AAV-eGFP was loaded into an alginate gel and allowed to diffuse over 9 days. Transduction was too low to be practical and the dose was abandoned. **B.** Temperature of media was checked after various times of ultrasonication to determine how long ultrasound could be used before overheating the media. **C.** Collagen-GAG scaffolds were ultrasonicated and conditioned media was added to HEK293T cells. Ultrasound-of the media alone (US-Cell Media) did not affect cell viability; however, sonication of the collagen-GAG scaffold did reduce viability, suggesting we should focus the ultrasound just on the pocketed regions of the scaffold (error bars = mean ± st dev; ANOVA with Tukey post hoc; * = $p<0.05$).

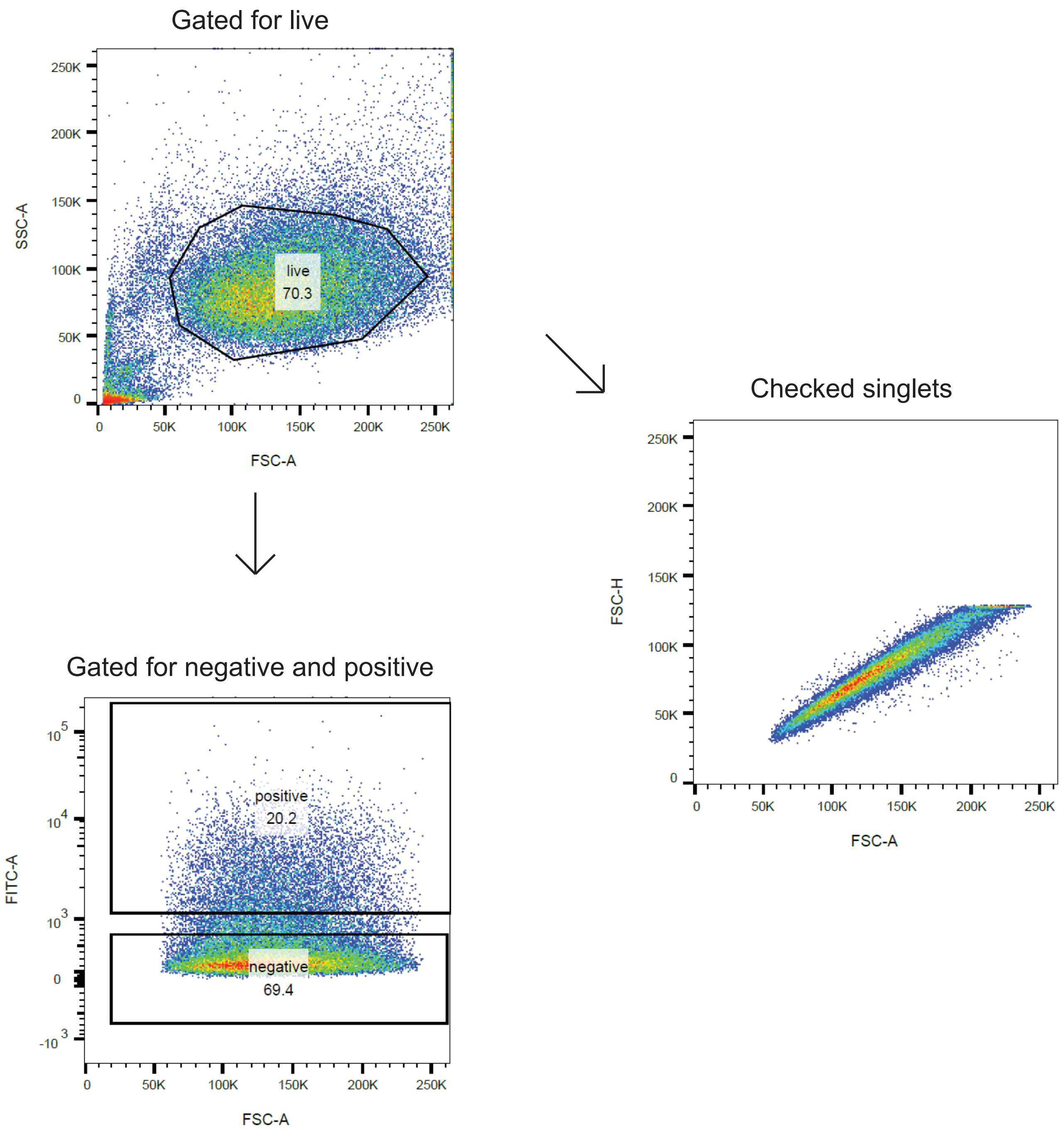


**Supplemental figure 3:** Example of flow cytometry analysis. First cells were gated on the SSC-A vs FSC-A axes for live cells. The live population gate was also used to confirm cells were in single suspension. Then that population was gated for negative (gate set by using negative control) and positive (gate set by including area above negative gate).

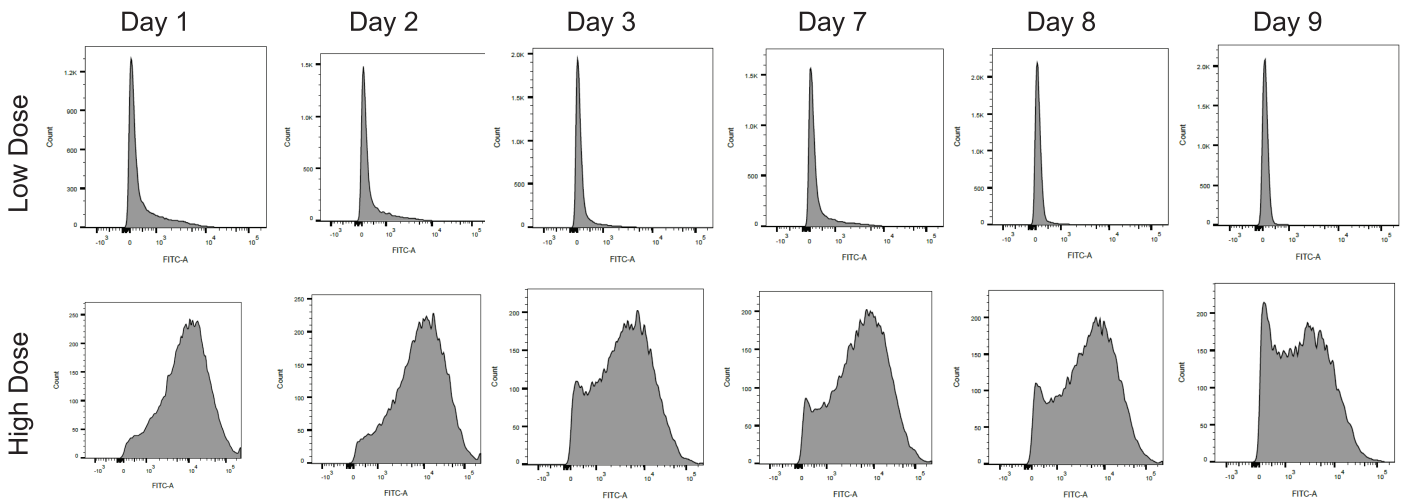


**Supplemental Figure 4:** Representative data of histograms showing fluorescent intensity. This is specifically from Figure 3E, and shows histograms after gating for live cells as shown in Supplemental Figure 3.

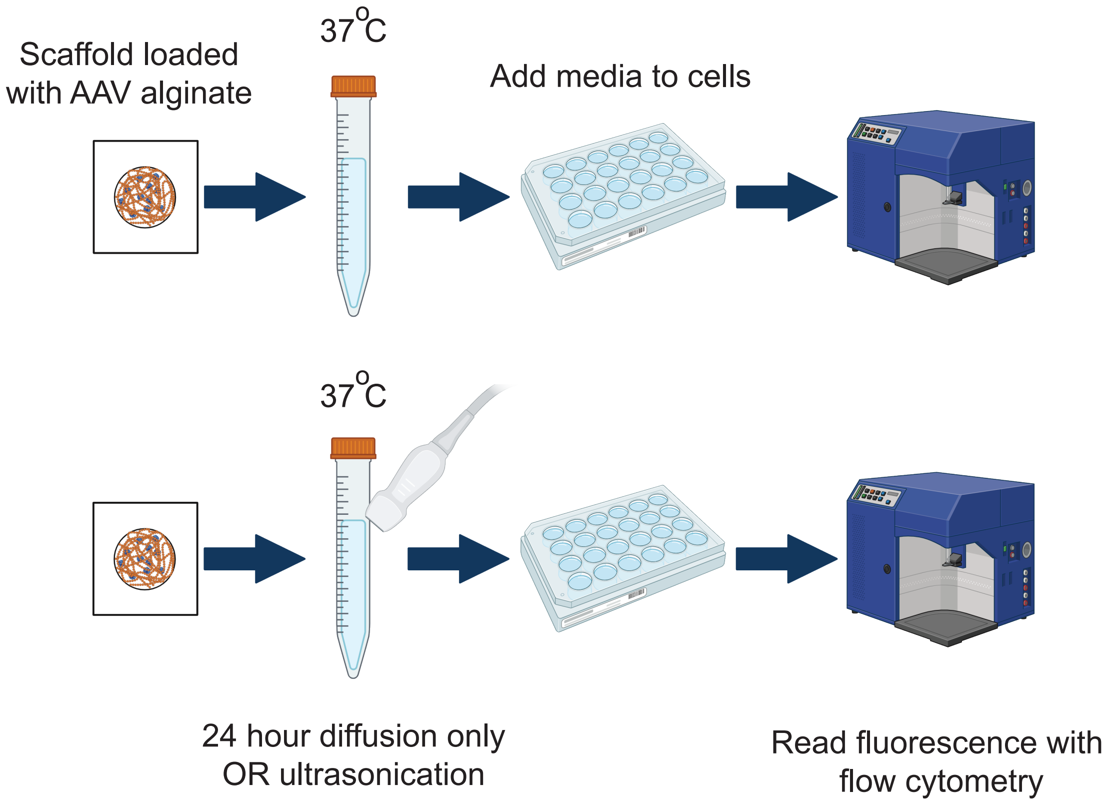


**Supplemental Figure 5:** Schematic showing work flow of AAV release experiments. First AAV loaded alginate was injected into the collagen-GAG scaffolds. Then media was added and AAV was allowed to diffuse for 24 hours. At the end of 24 hours ultrasound-treatment groups were ultrasonicated. Media was then collected and added to HEK293t cells. 2 days after addition cells were lifted and fluorescence was read via flow cytometry. For multi-day experiments media on the scaffolds was replaced when removed and scaffolds were stored at 37 $^{\circ}$ C.
